# Observations of anisotropic paramagnetic and diamagnetic susceptibility in the primate brain

**DOI:** 10.1101/2025.11.21.689628

**Authors:** Jingjia Chen, Dimitrios G. Gkotsoulias, Jenny E. Jaffe, Tobias Gräßle, Carsten Jäger, Zoro B. Goné Bi, Catherine Crockford, Roman Wittig, Harald E. Möller, Chunlei Liu

## Abstract

Bulk magnetic susceptibility in the brain white matter is known to be diamagnetic and anisotropic due to the ordered myelin lipids. While paramagnetic iron is widely present in the brain, it is typically not considered to contribute to anisotropy. Using experimental MRI and computational techniques, this study explores the competing contribution of diamagnetic and paramagnetic substances to the susceptibility anisotropy. Multi-echo gradient-echo imaging and diffusion-weighted imaging data from a paraformaldehyde-fixed *post-mortem* chimpanzee (*Pan troglodytes verus*) brain was analyzed. A computational method, DECOMPOSE-QSM, was used to separate paramagnetic susceptibility and diamagnetic susceptibility components. As expected, diamagnetic components showed significant anisotropy; unexpectedly, paramagnetic components also exhibited strong anisotropy in deep gray matter and parts of white matter. This may arise from the geometric arrangement of iron-rich cellular compartments, such as oligodendrocytes, astrocytes, and microglia, along nerve fibers. This method enables further exploration of tissue-specific contributions to susceptibility anisotropy.

## Introduction

Biological tissue’s bulk magnetic susceptibility (often labeled as χ) can be measured non-invasively with magnetic resonance imaging (MRI) employing a technique called quantitative susceptibility mapping (QSM) ^1–3^. QSM has been used as an MRI contrast for studying brain structure ^4,5^ and disease pathologies ^6–11^. Potential confounds in applications of QSM may result from the complex tissue composition and microstructure of an imaging voxel. In many brain regions, the measured susceptibility depends on the angle between the underlying structure and the direction of the static magnetic field of flux density **B**_0_ ^12–14^. Similar to the description of diffusion anisotropy, susceptibility anisotropy can be described by a second-order tensor. Using susceptibility measurements at multiple **B**_0_ orientations, the second-order susceptibility information can be characterized using the susceptibility tensor model, namely the method of susceptibility tensor imaging (STI) ^15^. It has been shown that the major contribution to susceptibility anisotropy in cerebral white matter (WM) is from the radially arranged myelin lipids concentrically wrapped around the axon. Therefore, STI is able to reflect the orientations of WM fiber tracts ^12,16–19^.

The coexistence of paramagnetic and diamagnetic susceptibility sources within a voxel affects susceptibility tensor quantification. Due to the strong contribution from the radially arranged myelin lipids to susceptibility anisotropy, most of the STI studies are focused on revealing WM fibrous structures, and the effect of paramagnetic contributions on the susceptibility anisotropy is typically ignored. However, a co-existing isotropic or mildly anisotropic paramagnetic susceptibility could, in a way, dilute the effect of the diamagnetic anisotropy. Moreover, the complex sub-voxel susceptibility mixture on the molecular level complicates investigations of potential contributions from paramagnetic susceptibility sources to susceptibility anisotropy.

There have been several efforts put into modeling and resolving the mixed susceptibility sources with different MRI protocols ^20–24^. Among those, the χ-separation method used a linear model to differentiate the contribution of paramagnetic and diamagnetic susceptibility sources to the rate of radiofrequency (RF)-reversible dephasing, 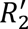, and bulk susceptibility, which requires measurements of both the transverse relaxation rate (*R*_2_ = 1⁄*T*_2_) and the effective transverse relaxation rate (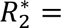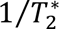) ^24^. Another method that only uses gradient-recalled echo (GRE) signals, without the need of *T*_2_data, namely DECOMPOSE-QSM ^20^ was proposed to achieve the same separation using a complex multi-exponential nonlinear signal model.

Here, we apply the DECOMPOSE-QSM method to a multi-orientation, multi-echo (ME)-GRE dataset of a *post-mortem* chimpanzee (*Pan troglodytes verus*) brain. We investigate the distinct anisotropic properties of diamagnetic and paramagnetic contributions to the bulk susceptibility. We hypothesize that the origin of the herein discovered paramagnetic components’ contribution to susceptibility anisotropy reflects the specific geometric arrangement of iron-rich cell types such as oligodendrocytes, astrocytes, and microglia.

## Results

### DECOMPOSE-STI improves spatial coherence of tensor element maps

Susceptibility tensors are calculated using multi-directional measurements of paramagnetic component susceptibility (PCS) and a diamagnetic component susceptibility (DCS) as well as standard QSM, referred to as PCS-STI, DCS-STI, and QSM-STI, respectively (Figure 1). The multi orientation QSM, PCS, and DCS maps used for STI calculation are shown in Supplementary Figure S1. For a representative axial slice, Figure 2A shows the diagonal elements of each tensor, and Figure 2B shows the off-diagonal tensor elements. The three principal eigenvalue maps of the corresponding slices are shown in Supplementary Figure S2. Overall, the PCS results align with the paramagnetic (bright) region in conventional QSM-STI, while the DCS results align with the diamagnetic (dark) regions in conventional QSM-STI. QSM-STI contrast shows the dominant susceptibility at each voxel, therefore, little diamagnetic anisotropy is revealed in deep gray matter (GM) and limited paramagnetic anisotropy is revealed in white matter. Tensor element maps from DECOMPOSE-STI, both PCS and DCS based, show spatially continuous and notable tissue susceptibility variations. For example, in the basal ganglia region, the paramagnetic contribution is dominant in conventional QSM-STI (i.e., positive sign of the QSM-based STI), while DCS-STI shows the presence of an additional non-zero diamagnetic susceptibility contribution. In the off-diagonal element maps, QSM-STI contains slightly more prominent streaking artifacts compared to the off-diagonal elements of PCS-STI and DCS-STI.

**Figure 1:**
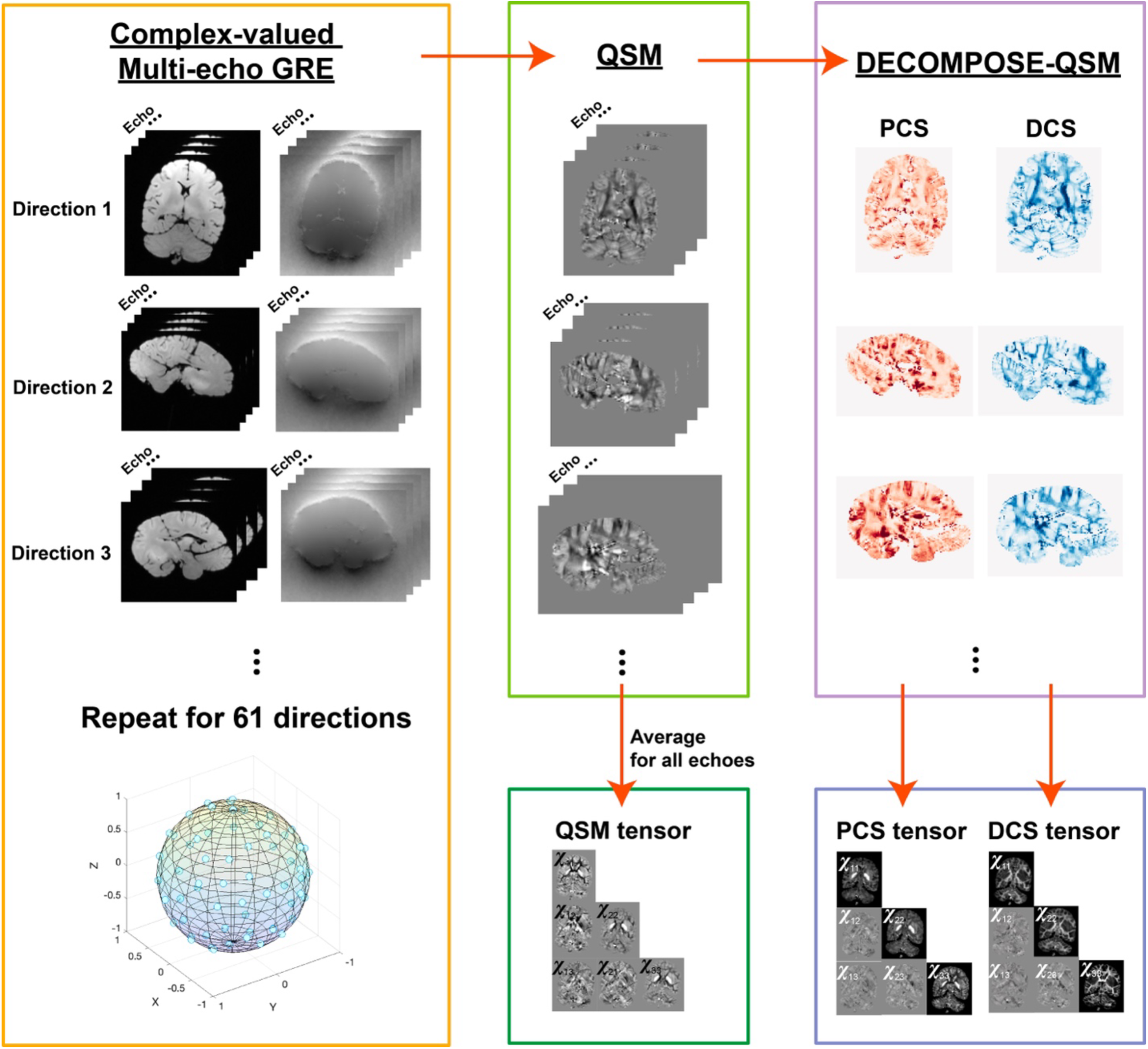
Schematic processing pipeline for obtaining susceptibility tensors. Complex-valued ME-GRE data of each direction is used to reconstruct susceptibility maps echo by echo. Then, the echo-dependent maps with the original ME magnitude images are used for DECOMPOSE-QSM producing PCS and DCS maps of each orientation. 61 directions of the echo-averaged QSM, PCS, and DCS are then used to calculate corresponding susceptibility tensors. The B_0_ field orientation is calculated through the rotation component of the rigid body transformation from each volume to the reference (the first direction) volume.

**Figure 2:**
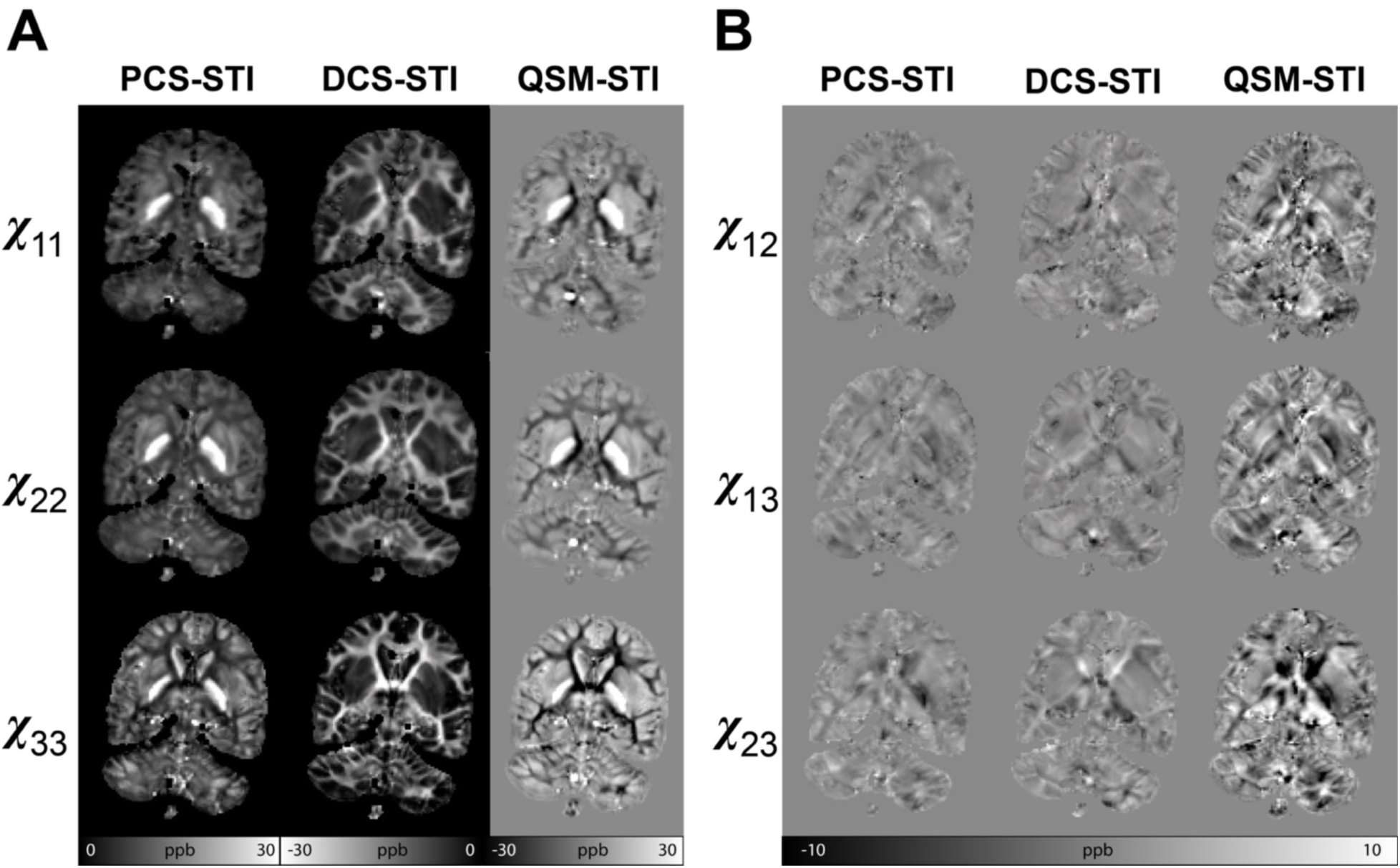
Tensor element maps of PCS, DCS, and QSM-based susceptibility tensors. (A) The three diagonal elements of the susceptibility tensors obtained by different methods. The PCS-based tensor has only positive values corresponding to paramagnetic susceptibility, and the DCS-based tensor has only negative values corresponding to diamagnetic susceptibility. The conventional QSM-based tensors have both positive and negative values, with positive values reflecting an overall paramagnetic average and negative values reflecting an overall diamagnetic average voxel composition. The DECOMPOSE-based method shows continuous tissue susceptibility changes while the QSM-based STI maps show sharp transitions between positive and negative susceptibility values. (B) The off-diagonal elements of susceptibility tensors. The QSM-based maps contain slightly more prominent streaking artifacts. The DCS-based tensor elements are shown in inverse contrast for better visualization.

### DTI fractional anisotropy (FA) and susceptibility anisotropy (SA)

Figure 3 shows the DTI-FA map and SA maps calculated using PCS-STI, DCS-STI, and QSM-STI (see Eqs. 4 and 5). The DCS-based SA is the most similar to the DTI-FA map, with both maps delineating the major WM structures (e.g., the corpus callosum, the internal capsule, the anterior corona radiata). SA maps of the QSM-STI appear to be a composite of SA maps from PCS-STI and DCS-STI. Due to the composition appearance, the QSM-based SA map does not delineate anatomical structures as well as the separated SA maps obtained from the PCS-STI or DCS-STI. As shown in Supplementary Figure S3, while the FA threshold level increases, the mask concentrates at the major WM tracts where the orientation of the tract is known to be more coherent. The center of the DCS-based SA distribution shifts towards higher values as the FA threshold increases, while the center of the PCS-based SA distribution remains at around 0.003 ppm.

**Figure 3:**
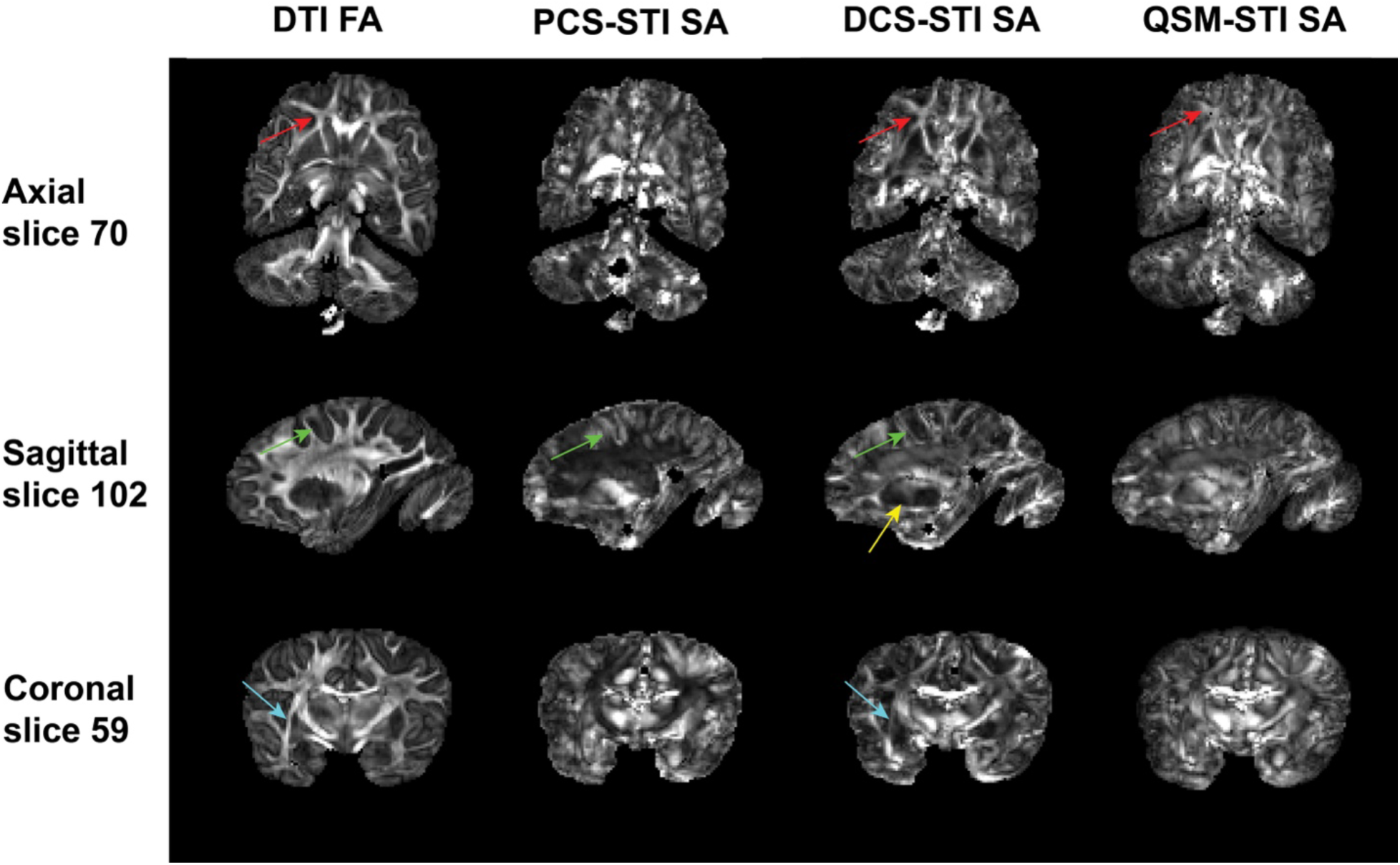
Diffusion tensor-based fractional anisotropy (FA) and susceptibility anisotropy (SA) comparison. DTI-FA and PCS-, DCS- and QSM-based SA of three representative slices are compared. Despite the differences in the origin of the two types of anisotropies, the DCS-based SA map looks the most similar to the DTI-FA map. The major WM tracts are bright indicating high structural and susceptibility anisotropy. The PCS highlights the SA in deep gray matter indicating the existence of underlying anisotropic susceptibility arrangements, which are not captured in DTI-based anisotropy measurements. QSM-based SA overall appears to be the composite of PCS- and DCS-based anisotropy. Regions that show strong differences between PCS- and DCS-based anisotropy are highlighted by colored arrows.

### FA-weighted primary eigenvector maps of diffusion tensors and susceptibility tensors

Figure 4 shows primary eigenvector maps obtained from the different tensors using an RGB color-coding. The DTI-FA map is used to weight the RGB map such that there is a focus on major WM tracts that can be identified by DTI. Overall, primary eigenvectors of all tensors align well in major fiber bundles, for example, the dominant red color from the body section of the corpus callosum, the purple/blue color of the corticospinal tracts, the green color of the superior longitudinal fasciculus and the inferior longitudinal fasciculus (Figure 4). The angle differences comparing eigenvectors of STI and DTI in the white matter region are shown in Supplementary Figure S4. Similar to the observations from the SA maps, the QSM-STI primary eigenvector seems to be the mixture of the primary eigenvectors obtained from PCS-STI and DCS-STI. Within the external capsule, DCS-STI is the only susceptibility-based tensor that recovers a similar fiber-direction pattern as obtained with DTI.

**Figure 4:**
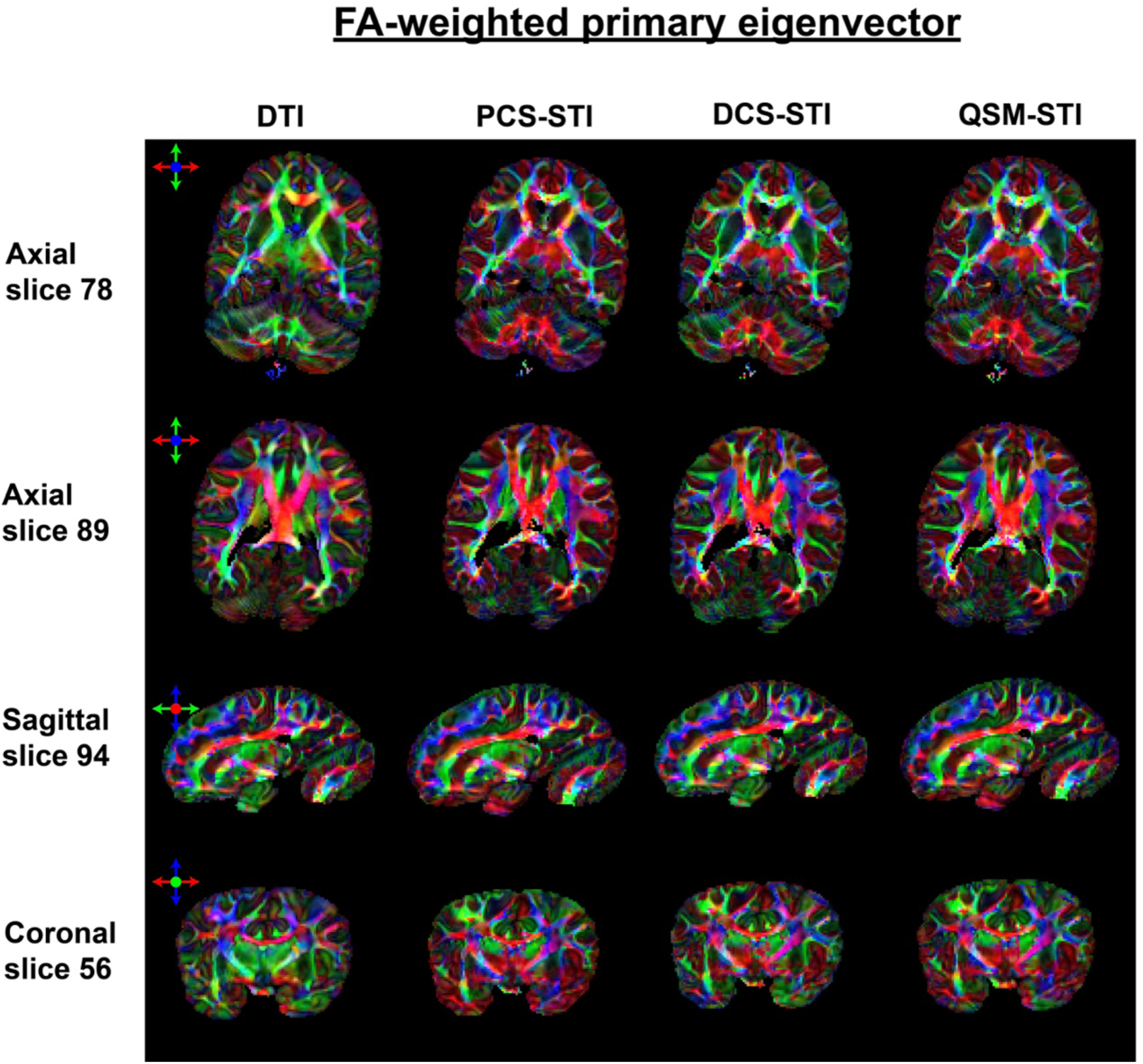
RGB Color-coded FA-weighted primary eigenvector maps obtained with DTI, PCS-STI, DCS-STI, and QSM-STI (red: left-right, green: anterior-posterior, blue: superior-inferior direction). Overall, vectors within major WM tracts (e.g., corpus callosum, corticospinal tracts, etc.) are showing alignments for all four types of tensors. The external capsule in the DCS-based susceptibility eigenvector map appears to be closest to the DTI-based eigenvector result.

### White matter structures

Figure 5 presents a few examples of zoomed-in views of WM structures and the corresponding SA maps are presented in Supplementary Figure S5. Figure 5A zooms in on the level of the basal ganglia. The red structure (highlighted by the yellow line) in between putamen and globus pallidus stands out in PCS-STI, DCS-STI, and QSM-STI eigenvector maps, coinciding with the lateral medullary lamina. In comparison, DTI only shows a hint of this structure in the corresponding area. In Figure 5B, major figure bundles such as corpus callosum are revealed clearly through all maps. The external capsule (to the left of the yellow line) stands out on the DCS-STI eigenvector maps, matching the appearance on the DTI result. However, the PCS-STI and QSM-STI are not able to depict this thin structure at the native 1mm isotropic resolution. The zoomed-in view of Figure 5C depicts a cortical region and adjacent superficial WM. The DCS-STI shows curving structures following the gyrus whereas QSM-STI, PCS-STI and DTI in this region do not seem to have high alignments in terms of the eigenvectors.

**Figure 5:**
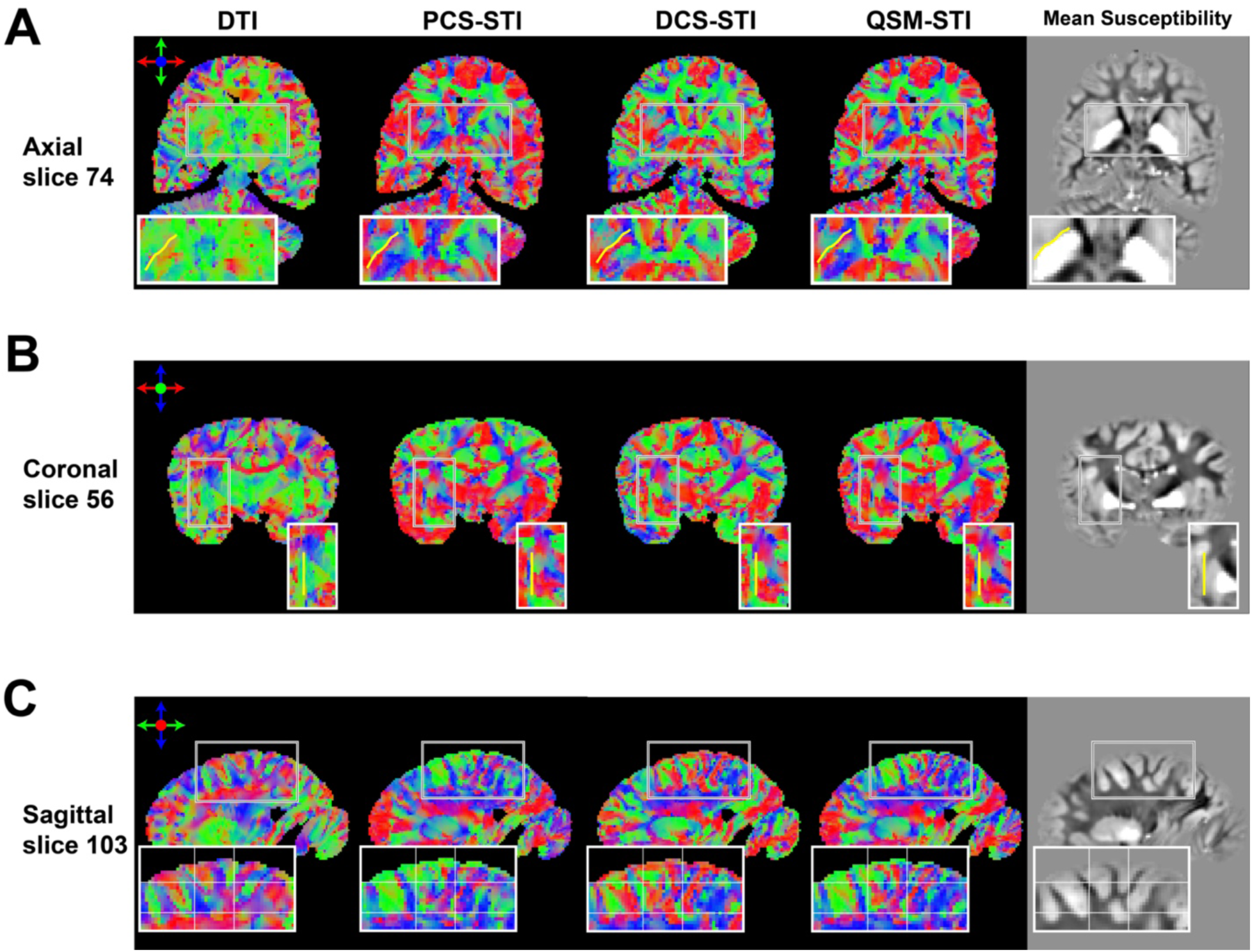
Color-coded primary eigenvector maps of DTI, PCS-STI, DCS-STI, QSM-STI with zoomed-in views into three WM regions. (A) Magnified region highlighting the lateral medullary lamina (yellow line). (B) Magnified region highlighting the external capsule (yellow line). (C) Magnified region in superficial WM. A grid is added to guide a visual comparison.

### Deep gray matter structures

The non-weighted and RGB color-coded primary eigenvector maps with zoomed-in views of deep GM regions are shown in Figure 6. The corresponding SA maps are shown in Supplementary Figure S6. Overall, the susceptibility tensors reveal more orientation details in the deep GM, while the diffusion tensor shows subtle contrast in terms of revealing structure orientations. Figure 6A shows a zoomed-in view of the substantial nigra (SN). While DTI primary eigenvector map is homogeneous throughout the SN and its surrounding area, suggesting that the anisotropy information at this area is unreliable. The PCS-based primary eigenvector shows the most coherent directions within the SN region, while DCS- and QSM-based eigenvectors show some spatial variations within the region as well, potentially suggesting underlying structures within the SN region. A zoomed-in view of the basal ganglia region is shown in Figure 6B. The PCS-based eigenvector shows distinct subregional orientations within the putamen and the globus pallidus. The eigenvector from DCS-STI shows a similar trend, but the contrast is less prominent and continuous compared to the PCS-based eigenvector map. Figure 6C shows a zoomed-in view of the thalamus. The STI-based eigenvector and SA maps show groups of coherent structures that resemble thalamic nuclei, while the DTI eigenvector maps only exhibit uniform patches within the thalamus.

**Figure 6:**
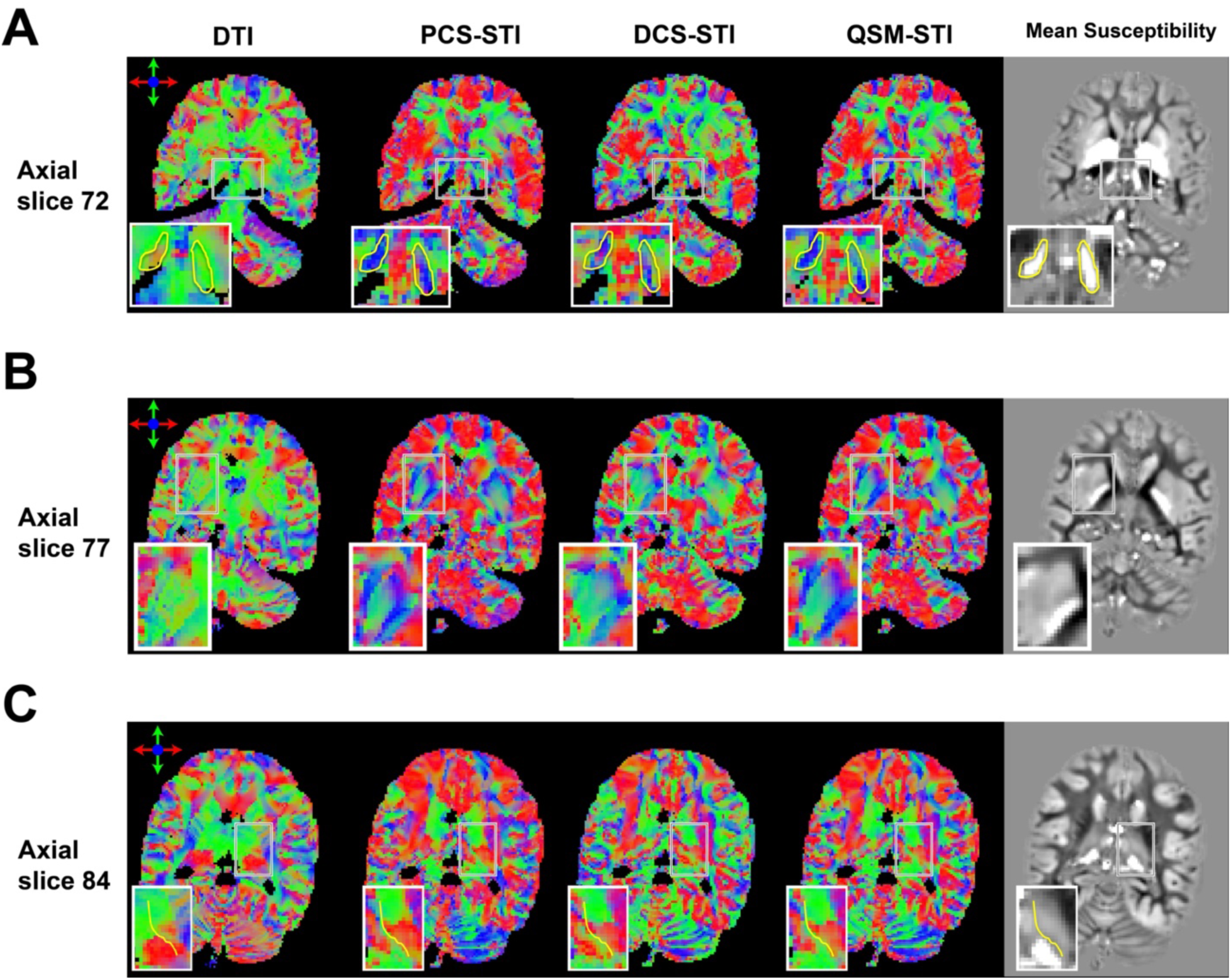
Color-coded primary eigenvector maps of DTI, PCS-STI, DCS-STI and QSM-STI, with a zoomed-in view of deep GM regions. (A) Magnified region around the substantia nigra (SN) with highlighted SN boundary by yellow lines. Within this boundary, the PCS-STI shows the most coherent eigenvector direction. (B) Magnified region at the level of the basal ganglia. The PCS-STI shows a continuous eigenvector direction distribution in putamen and globus pallidum. (C) Magnified region at the level of the thalamus with the internal medullary lamina indicated by yellow lines.

## Discussion

### Paramagnetic susceptibility anisotropy

Susceptibility anisotropy in cerebral WM was observed in multiple previous STI studies^12,15,17,25–27^. The radially aligned multi-sheath double-layered myelin lipid molecules are considered to be the main sources of the macroscopic susceptibility anisotropy in the brain (Wharton and Bowtell, 2012). According to the radially arranged lipid shell model, which is focused on the diamagnetic nature of the myelin lipids, the strongest negative values of the observed orientation-dependent susceptibility are obtained when **B**_0_ is parallel to the axon’s long axis. Therefore, the primary eigenvector of the susceptibility tensor reveals the fiber orientation in such WM regions. This model is largely correct when the underlying microstructure is dominated by well-aligned myelinated fiber bundles. However, these studies fall short in considering separate contributions to anisotropy from diamagnetic and paramagnetic susceptibility. With additional decomposition into diamagnetic and paramagnetic contributions in the current work, this notion is verified by the obtained DCS-STI results. Additionally, we observed that the PCS susceptibility also exhibits orientation-dependent character in WM, but more prominently in the deep GM, where paramagnetic sources are generally more abundant^28^.

Though the source of the magnetic susceptibility anisotropy is considered to occur from the radial arrangement of the diamagnetic myelin lipid around axons^12^, certain other geometric arrangements of microscopic structures may also introduce anisotropy of the local bulk susceptibility observed at the level of an imaging voxel (on a mm^3^ spatial scale). According to the previously proposed hollow cylinder model of the myelin sheath^13,14^, at a molecular level, both isotropic-only and anisotropic-only susceptibility molecular configurations may produce angle-dependent frequency maps. In WM, considering the diamagnetic nature of the dominant myelin lipids, it is the anisotropic-only susceptibility molecular configuration that is expected to better fit *in vivo* measurements. As for the isotropic-only susceptibility molecules, if the geometry of the molecule arrangement still follows hollow cylinder shape, they still exhibit magnetic anisotropy when **B**_0_ is perpendicular to the axon.

Therefore, we hypothesize that the observed paramagnetic susceptibility anisotropy originates from paramagnetic sources located around myelinated axons or intra-axonally. These paramagnetic sources can be iron (in the form of ferritin) in oligodendrocyte cell bodies, astrocytes, and microglia, which is consistent with the known arrangement of clusters of glial cells in short rows parallel to the axons they support ^29–31^. A numerical simulation (Supplementary Figure S7) of the PCS anisotropy with various **B**_0_ angles shows that when paramagnetic particles are distributed around an axon in a cylindrical fashion, the apparent susceptibility will show anisotropic behaviour with the highest apparent susceptibility value when **B**_0_ is aligned with the axon axis. This suggests that susceptibility caused by radially arranged myelin lipids also from cylindrically distributed paramagnetic compounds (such as iron) both show the maximum signed value when **B**_0_ is aligned with the axon axis. In other words, the primary eigenvectors from PCS-STI and DCS-STI should be parallel. In Figures 4–6, major WM structures, such as corpus callosum and corticospinal tracts, show a high agreement of the primary eigenvectors, which supports this hypothesis.

Previously, a histology-based technique was used to reveal the WM fiber structures in the brain ^31^. In this work, Nissl staining in multiple human brain sample slices was used to reconstruct the structure tensor and glial row orientation density function based exactly on the glial organization around myelinated axons. The orientation map derived from the Nissl-based structure tensor aligned very well with previously published fiber orientation maps based on *post-mortem* polarized light imaging (PLI). This once again supports the assumption of an anisotropy contribution from iron-bearing cellular structures. It is worth noting that the iron distribution is still less ordered than the myelin structure. Therefore, in the WM, the SA from DCS-STI is higher than the SA from PCS-STI (Figure 3).

### Paramagnetic and diamagnetic susceptibility anisotropies coexist

We observe that both paramagnetic iron-rich tissue components and diamagnetic compounds may contribute to magnetic susceptibility anisotropy depending on the specific tissue composition and their geometric arrangement. From Figures 3, 5, and 6, QSM-based STI appears to show the summation effect of the DCS-STI and PCS-STI. In regions with high paramagnetic susceptibility, the SA and the primary eigenvector direction obtained with QSM-STI are close to those obtained with PCS-STI (e.g., deep GM). In regions with high diamagnetic susceptibility, the SA and the primary eigenvector direction of QSM-STI are close to that in DCS-STI (e.g., WM tracts). The conventional QSM-STI image interpretation can be problematic due to simultaneous contributions from diamagnetic and paramagnetic compounds since iron and myelin both cause opposite frequency shifts. If STI is based on separated susceptibility contributions, this could provide additional information on the paramagnetic and diamagnetic compounds as well as their anisotropic distribution in a tissue. In the current dataset, DCS-STI shows cleaner directionality and is closer to the DTI results compared to those from QSM-STI and PCS-STI, especially for the external capsule, superior longitudinal fasciculus and inferior longitudinal fasciculus. In such regions, QSM yields values close to zero, whereas after the decomposition, separating the PCS-STI component acts as a ‘denoising’ process, by removing the non-diamagnetic susceptibility anisotropy, which results in a more coherent eigenvector estimation of DCS-STI. Similarly, in the iron-rich basal ganglia, PCS-STI should provide more reliable information about the underlying composition and microstructure.

### DCS anisotropy is similar to diffusion anisotropy in white matter

Overall, DCS-based SA appears the most similar to the diffusion FA compared to PCS- and QSM-based SA in WM (Figure 3 and Figures S3, S4 and S5), presumably because the are both sensitive to the same aspects of the tissue composition and microstructure. The FA-weighted color-coded primary eigenvector maps from DCS-STI also resemble the DTI-based primary eigenvector maps better (Figure 4). In major WM fiber bundles, SA is mostly due to the radially arranged myelin lipids wrapping around the axon, in line with the high agreement of the DCS-based SA and FA. However, the physical mechanisms underlying FA and SA are different. The FA reflects the anisotropic hindrance of water diffusion while SA describes the magnetization differences applying **B**_0_ at different directions. Similar (and potentially complementary) to DTI, susceptibility anisotropy yields information about the tissue microstructure. For example, in previous research on the kidney^32^, renal tubules were revealed in both the inner medulla and outer medulla using STI, whereas DTI only showed clear fibrous structures within the inner medulla.

### PCS and DCS anisotropies in and around deep gray matter

In WM, the major contributions to diamagnetic and paramagnetic susceptibility anisotropy are the myelin sheath and the local order of surrounding cell bodies caused by the presence of fiber bundles, respectively^13,14,33^. Hence, these eigenvectors align largely with the diffusion-based eigenvector estimations. In deep GM, the tissue’s physical structure becomes more complex, and the diffusion tensor model may not perform as well as in the major fiber bundles in terms of providing information about the tissue substructure. STI probes the tissue’s magnetic properties and the microstructural arrangement of magnetized compounds. In Figure 6A, the STI-based fiber directions in substantia nigra are primarily along the superior-inferior direction, which agrees with previous human brain high-resolution STI at 7T^25^. In Figure 5A, a red color-labeled structure (indicating primarily left-right direction) between putamen and globus pallidus is clearly visible in all susceptibility-based eigenvector maps. It could correspond to the thin WM lateral medullary lamina connecting the putamen and globus pallidus. After separation of the susceptibility sources, the diamagnetic WM contribution to the anisotropy becomes purer compared to the simple QSM-STI result. Therefore, the lateral medullary lamina sandwiched in between two predominantly paramagnetic nuclei become clearer and cleaner on DCS-based eigenvector maps. In Figure 6B, the complex directional information in putamen and globus pallidus is recovered using all STI measures. However, since the iron-rich basal ganglia are paramagnetic, and the largest SA is obtained with PCS-STI, and the corresponding eigenvector map is cleaner compared to the DCS-STI result.

### Limitations and future work

QSM-based STI and DECOMPOSE-STI show promising results and potential in revealing tissue microstructure in addition to or complementary to DTI. However, some STI (of all types) results do not align and are even perpendicular to the diffusion tensor, especially in cortical regions (Figure 5C). This could be due to several reasons: 1) increasing inaccuracy of the susceptibility estimation in voxels closer to the surface of the brain; 2) the tissue microstructure may be too complex for the simple single-exponential STI and time-independent DTI models; 3) the physical origins of the diffusion anisotropy and susceptibility anisotropy are fundamentally different. A previous study ^34^ has found that diffusion FA did not match the spatial distribution of myelin in the GM but agreed better with Nissl-stained tissue anisotropy. In this study, the DECPMPOSE-STI is achieved through the incorporation of DECOMPOSE-QSM, a model based on isotropic spherical susceptibility sources, into the susceptibility tensor model. While isotropic spheres can be viewed as a basis for generating the field perturbation of susceptibility sources of other geometries, more sophisticated tissue microstructure diffusion models and multi-compartment susceptibility models are needed to coordinate different underlying effects that may contribute to the tissue anisotropy measurements.

The paramagnetic susceptibility anisotropy was observed in this study and hypothesized to be related to the distribution of iron. To further investigate or validate this assumption about the origin of paramagnetic susceptibility anisotropy, iron-washed brain tissue samples could be deployed and followed by the same imaging experiment setups. Additionally, higher-resolution datasets (of both STI and DTI), PLI on sample slices, or the Nissl staining-based tensor reconstruction could provide additional information and potential validation.

The QSM-based SA and eigenvector maps appear to be a combination of the PCS- and DCS-based maps. At visual comparison, the DCS-STI seems more reliable when the tissue is diamagnetic and the DSC-based SA is higher. The effect is likewise for PCS-STI. For regions with higher content of paramagnetic susceptibility sources, the PCS-STI is more reliable in revealing the tissue anisotropy. A method to produce spatially informed combinations of PCS-STI and DCS-STI eigenvector maps could potentially lead to more informative maps of tissue orientational characteristics

In this work, no susceptibility value referencing was done to the QSM maps. The sample was from a wild animal that had died of natural causes at the age of 45 years and had no known brain pathology, so we hypothesize that the mean susceptibility of the whole brain should be around zero ^35,36^. However, when investigating a brain with pathologies or tissue with a non-balanced amount of diamagnetic/paramagnetic content, a sophisticated referencing procedure is needed prior to computing the DECOMPOSE-QSM, e.g., usage of an external reference.

Since the introduction of STI, many reconstruction advances have been proposed ^25,37–41^ to improve the reconstruction robustness as well as to obtain reliable tensors with as few orientations as possible. Those methods could be integrated with the susceptibility source separation method to extract more reliable information on the tissue susceptibility properties.

In summary, DECOMPOSE-STI, a DECOMPOSE-QSM preprocessed susceptibility tensor reconstruction, is introduced in this study. This proof-of-concept initial demonstration shows that the susceptibility source separation for susceptibility tensors could provide complementary information on tissue properties beyond the original STI approach and in addition to DTI. We have observed the existence of paramagnetic susceptibility anisotropy in a *post-mortem* brain sample without dissection of the brain, and we hypothesize that this is due to the arrangement of iron-bearing molecules (such as ferritin in glial cells) along the myelinated axons. Further, we show that separated susceptibility tensors could help investigate the tissue type contribution to the susceptibility anisotropy, and individual DCS and PCS tensor maps could provide a more coherent and reliable estimation of the underlying anisotropy.

## Materials and Methods

### Susceptibility tensor imaging in the lab frame

For tissue with anisotropic magnetic susceptibility, the susceptibility is no longer described by a scalar but by a second-order tensor. The MR field perturbation caused by an anisotropic magnetic susceptibility tensor 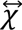 can be written as follows ^12,15^:

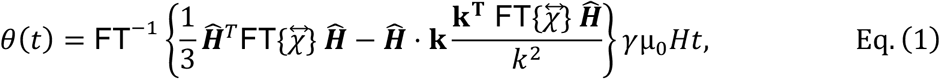

where *θ* is the phase of GRE signal at a certain sampling echo time *t*, ***Ĥ*** is the unit vector representing the static field with the magnitude being *H*, **k** is the frequency space coordinate, 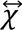 is the macroscopic susceptibility tensor described as a real-valued 3×3 matrix, *γ* is the gyromagnetic ratio, and FT means the Fourier transform operation.

The relation in Eq. (1) is defined in the subject reference frame. It can also be written in the lab frame (i.e., the frame defined by the scanner’s magnetic field) as

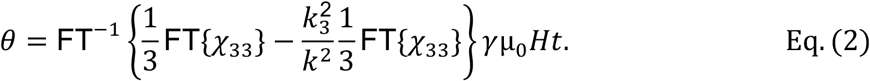

According to Eq. (2), the rotation of the sample with respect to the **B**_0_ direction (*z*-axis) corresponds to a rotation of the susceptibility tensor. Specifically, the *χ*_33_ tensor component (‘measured susceptibility’), which has been suggested as an STI-based estimation of a scalar susceptibility (Langkammer et al., 2018), is given by *χ*_33_ = ***R***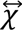***R***^*T*^. ***R*** is the rotation component of the transformation matrix from each direction to the reference direction.

### DECOMPOSE-QSM

The DECOMPOSE-QSM model aims to resolve the magnetic susceptibility sub-voxel mixture situation using the following multi-echo (ME) GRE signal model (Chen et al., 2021):

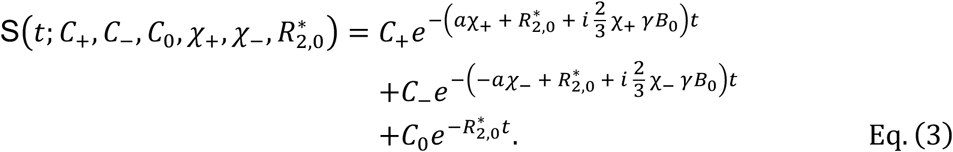

In Eq. (3), *C*_+_, *C*_–_ and *C*_0_ indicate the concentrations of the corresponding paramagnetic/diamagnetic/neutral (susceptibility reference; *χ* _0_ ≡ 0) components, *χ* _+/–_ are the volume susceptibilities of paramagnetic/diamagnetic components, 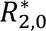 is the effective transverse relaxation rate, and *a* is a precalculated coefficient, which takes value of 323.5 Hz/ppm in a 3T MR system (Chen et al., 2021). The quantification metrics being used are a Paramagnetic Component Susceptibility (PCS) and a Diamagnetic Component Susceptibility (DCS), which are computed based on the signal model as follows:

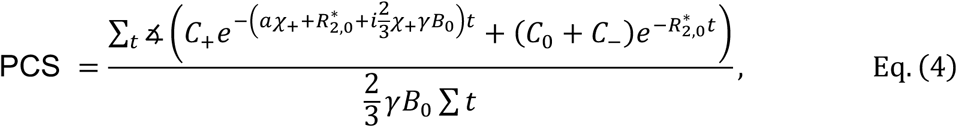

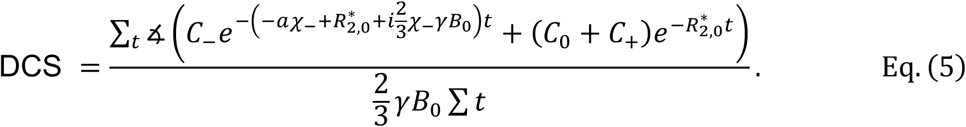

The rational for introducing PCS and DCS is that we want to re-create the voxel signal by considering only the paramagnetic or diamagnetic susceptibility sources embedded in the neutral component so that the resulting effective bulk susceptibility will be in comparable measures with the conventional QSM values.

### Sample preparation

The brain sample used in this experiment originates from a wild adult male chimpanzee (*Pan troglodytes verus*), who died from natural causes. The whole brain was extracted under strict biosafety, as detailed elsewhere^42^ within an acceptable *post-mortem* interval and preserved in 4% paraformaldehyde (PFA). After six months of fixation, prior to imaging, the PFA was washed out in phosphate-buffered saline (PBS) at pH 7.4 for 24 days. During imaging, the brain was submerged in a proton-free solution (Fomblin®; Solvay Solexis, Bollate, Italy). The sample was enclosed in a custom-designed 3D-printed container that supports re-orientations within the scanner head coil. The dataset was previously used and reported, along with all relevant details of ethics approval, extraction, preservation, and quality of the tissue as well as the measurements^16^.

### MRI acquisition

The inverse problem of the STI model is intrinsically ill-posed when the range of angles is limited. Various approaches^25,37–41,43^ have been proposed to improve the robustness of reconstructing susceptibility tensors while mitigating artifacts. In this study, since a *post-mortem* sample was used, images are acquired at a large number of angles.

A 3T MAGNETOM Skyra Connectom A (Siemens Healthineers, Erlangen, Germany) with a maximum gradient strength of 300 mT/m was used for MRI as detailed elsewhere.^16^ To record the susceptibility anisotropy, 61 unique directions of 3D ME-GRE (12 echoes) volumetric images were acquired with the following parameters: echo times TE_1_/TE_2_/ΔTE/TE_12_ = 3.54/6.98/3.75/44.48 ms, repetition time TR = 50 ms, and 1 mm isotropic native resolution. Additionally, diffusion tensor imaging (DTI) was also performed with 60 gradient directions and six non-diffusion-weighted images with a ME segmented 3D echo planar imaging (EPI) technique ^44^ and 1mm isotropic resolution, TR = 10.4 s, and b = 5000 s/mm^2^.

### Data processing and susceptibility tensor calculation

The data processing pipeline is illustrated in Figure 1. The complex-valued ME-GRE images from 32 individual coil elements were combined and processed using STISuite (https://people.eecs.berkeley.edu/~chunlei.liu/software.html) for each direction. Briefly, the phase of each coil element is first unwrapped using a Laplacian-based method^45^. Then, the 3D image volume is recovered by the weighted sum of the magnitude and phase. The unwrapped phase is filtered by V-SHARP ^46^ with a spherical mean value filter radius of 12 mm. Susceptibility maps of each echo were calculated using the STAR-QSM algorithm ^47^ with 12 mm padding and **B**_0_ input of each imaging orientation. The echo-dependent QSM and magnitude volumetric images of each direction were input into DECOMPOSE-QSM ^20^ to produce PCS and DCS maps (Supplementary Figure S1). The multi-direction volumes were registered to a reference volume (the first direction) using a rigid-body 6-parameter model transformation in FSL ^48^. The effective **B**_0_ field orientation of each direction is calculated using the rotation component of the transformation matrix. The transformation matrix was applied to QSM, PCS, and DCS, respectively. QSM averaged along the echo dimension was used for STI reconstruction in the lab frame ^15^. Similarly, PCS and DCS were used to calculate PCS and DCS tensors in the lab frame.

Eigen decomposition was performed on the calculated tensors. The three principal susceptibilities χ_1_, χ_2_ and χ_3_ are numbered in descending order. Susceptibility anisotropy, defined as SA = χ_1_ − (χ_2_ + χ_3_)/2, was calculated using the eigenvalues of the tensors (the first eigenvalue χ_1_is the largest eigenvalue). The eigenvector corresponding to the largest eigenvalue χ_1_ indicates the primary susceptibility (or susceptibility component) direction. Similar to the color-coding of diffusion MRI, RGB colors are used to color code the direction of the primary eigenvector as follows: left-right is coded as red, anterior-posterior is coded as green, and superior-inferior is coded as blue.

Each volume of diffusion-weighted images is registered to the ME-GRE space using an affine transformation with ITK-SNAP (Version 3.8.0) ^49^. The diffusion-encoding gradient orientations are transformed using the rotation component of the transformation matrix. The diffusion tensor of the volume is calculated in the ME-GRE space. DTI fractional anisotropy (FA) was used to weigh the RGB color-coded primary eigenvector map for diffusion tensors and susceptibility tensors. The mean magnetic susceptibility (MMS) defined as the trace of the QSM-based tensor, MMS = (χ_1_ + χ_2_ + χ_3_)/3, was calculated and used to provide anatomical guidance for the color-coded primary eigenvectors of the QSM-, PCS-, and DCS-based susceptibility tensors.

## Acknowledgements

This work was supported in part by the Alzheimer’s Drug Discovery Foundation through grant GC-201810-2017383 and by the EU through grant H2020-MSCA-ITN-2018 (“INSPiRE-MED”). Research reported in this publication was in part supported by the National Institute of Aging of the National Institutes of Health under Award Number R01AG070826. The content is solely the responsibility of the authors and does not necessarily represent the official views of the National Institutes of Health.

We thank the Ministère de l’Enseignement Supérieur et de la Recherche Scientifique and the Ministère des Eaux et Fôrests in Côte d’Ivoire, and the Office Ivoirien des Parcs et Réserves for permitting the study. We are grateful to the Centre Suisse de Recherches Scientifiques en Côte d’Ivoire and the staff members of the Taï Chimpanzee Project for their support and to Angela D. Friederici, Nikolaus Weiskopf and other Evolution of Brain Connectivity (EBC) project organizers for kindly sharing the brain specimen.

