## Supplementary Figure for "Observations of anisotropic paramagnetic and diamagnetic susceptibility in the primate brain"

**Supplementary Figure S1**: QSM, PCS, and DCS results of all 61 orientations. Middle axial slices are shown as an illustration.

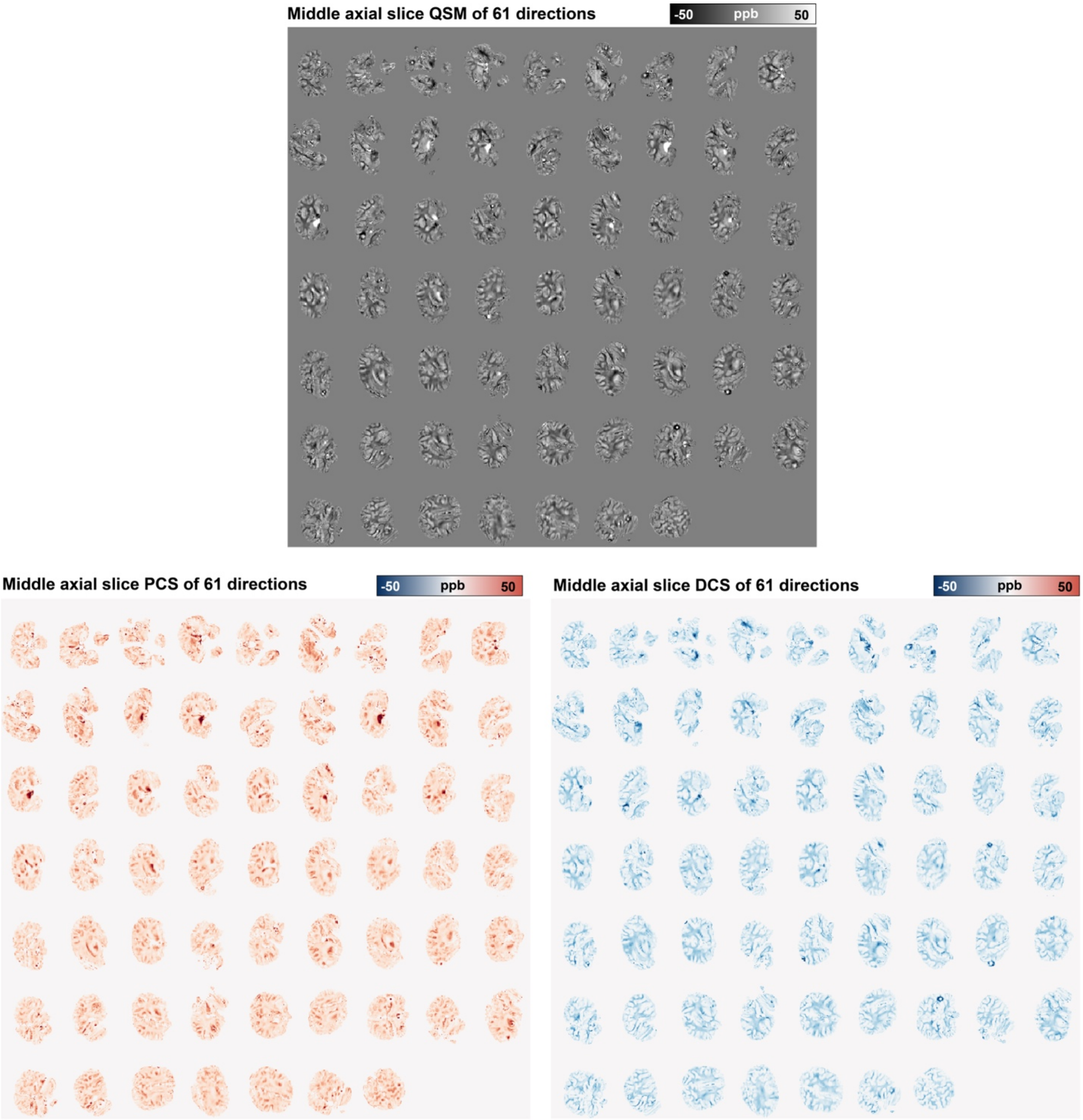

**Supplementary Figure S2**: Eigenvalues and the mean magnetic susceptibility (MMS) of PCS-, DCS-, and QSM-based susceptibility tensors.

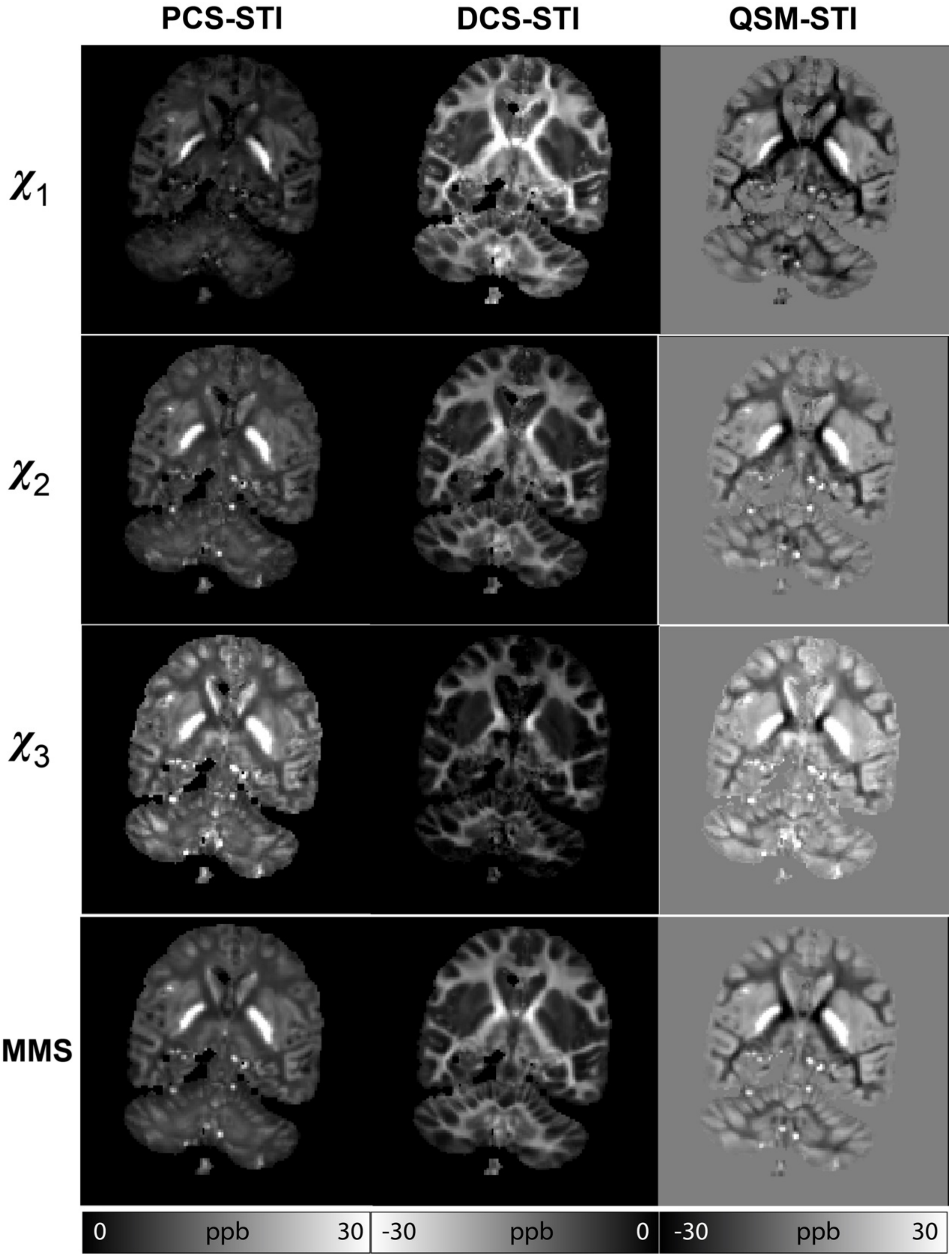

**Supplementary Figure S3**: SA distribution for different threshold levels of FA values. (A) With a higher FA threshold, the mask picks out regions with more coherent WM bundles. (B) The DCS-based SA distribution (histograms) shifts towards higher values while the PCS-based SA remains largely invariant as the FA threshold increases. The vertical green dashed line indicates the SA value at 0.003 ppm.

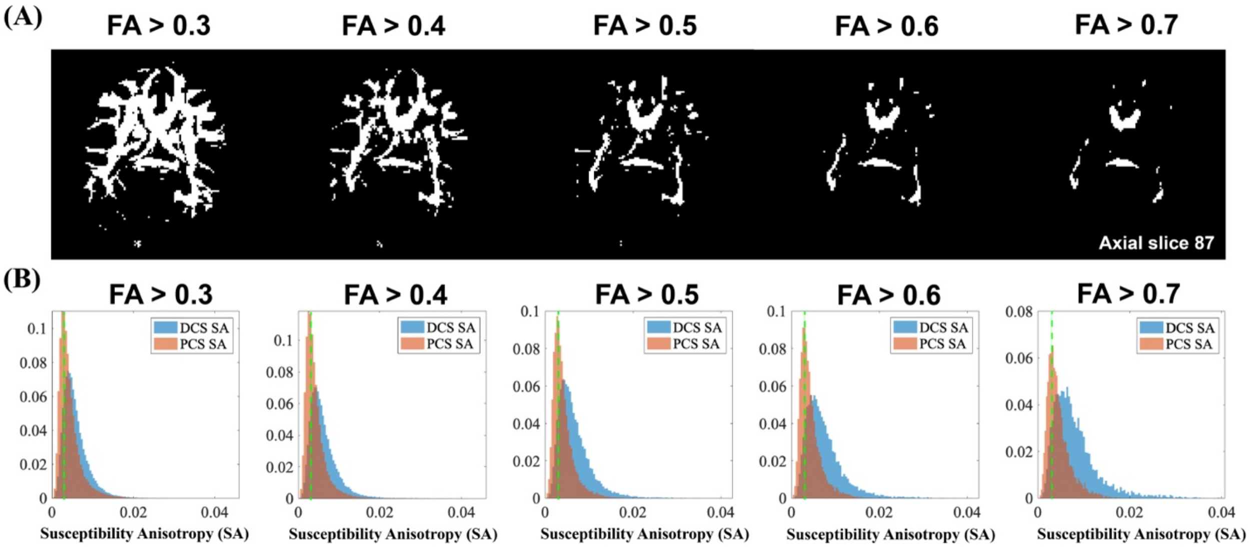

**Supplementary Figure S4**: Angle differences of primary eigenvectors between diffusion tensors and susceptibility tensors. A mask of FA > 0.3 is used to highlight the WM area. An angle of 0º means that the STI-based primary eigenvectors align exactly with the DTI-based primary eigenvectors. An angle of 90º means that the STI-based primary eigenvectors are perpendicular to the DTI-based primary eigenvectors. The green circle indicates the body part of the corpus callosum, where all types of susceptibility tensor eigenvectors align well with the diffusion tensor eigenvectors. The orange circle indicates the external capsule. Compared to the other two types of susceptibility tensors, the DCS-based tensor recovers a more similar direction as the diffusion tensor in this region.

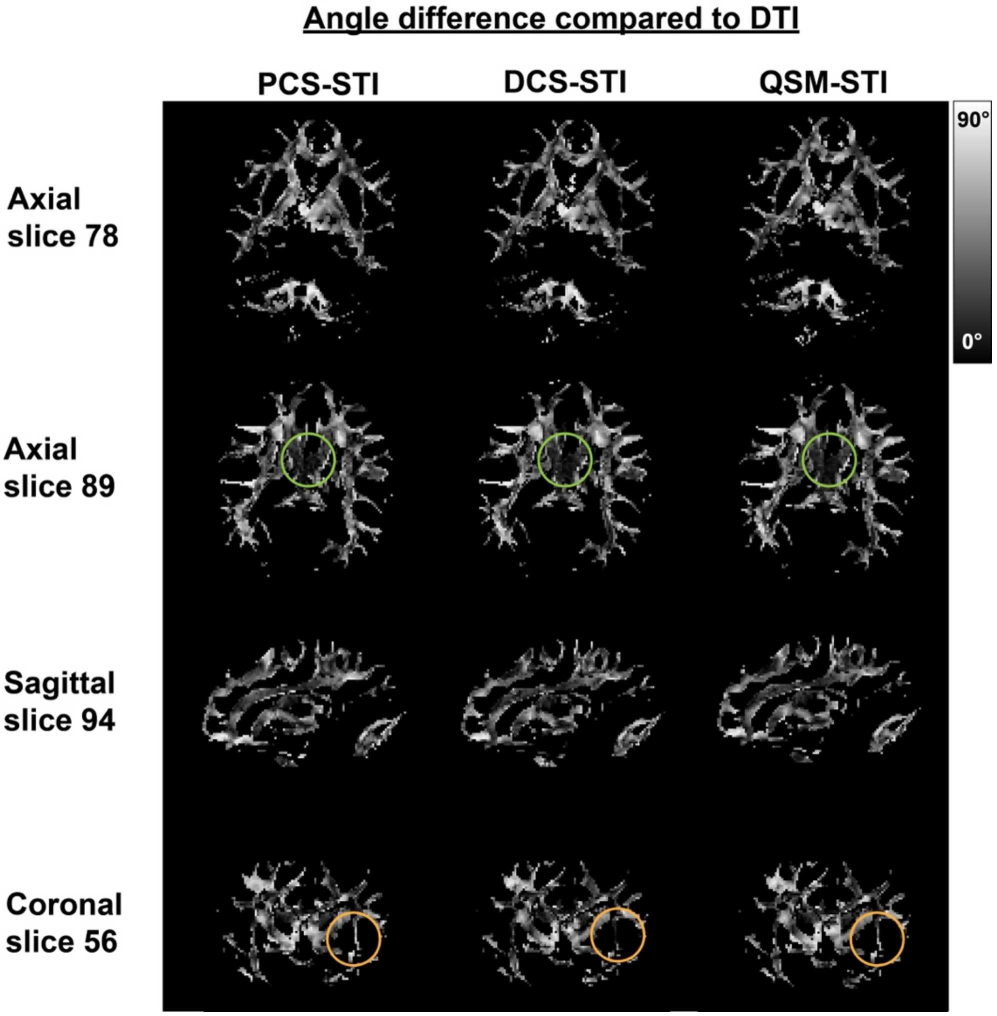

**Supplementary Figure S5**: FA and SA maps of the corresponding slices as shown in Figure 5. Bright contrast indicates high FA or SA.

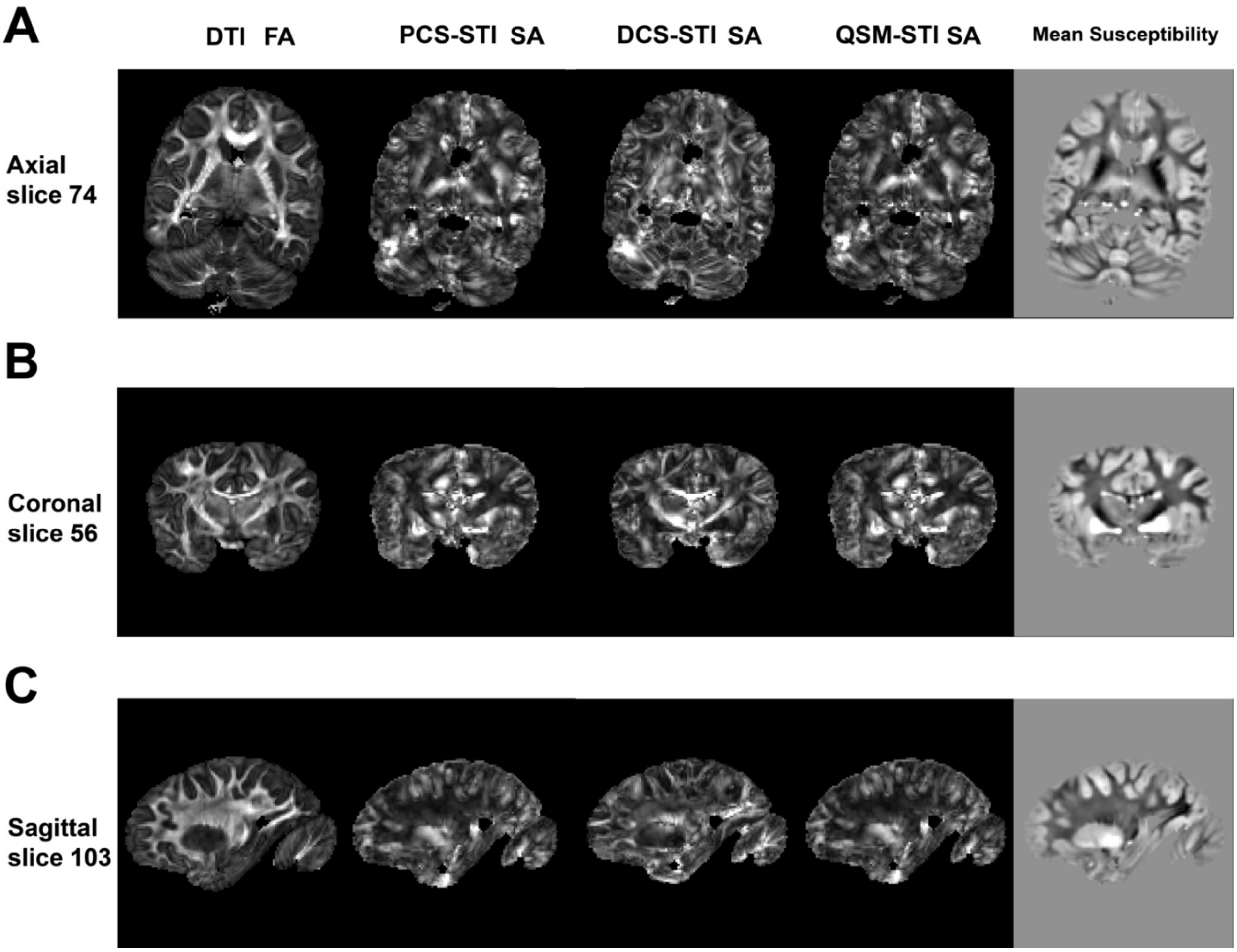

**Supplementary Figure S6**: FA and SA maps of the corresponding slices as shown in Figure 6. Bright contrast indicates high FA or SA.

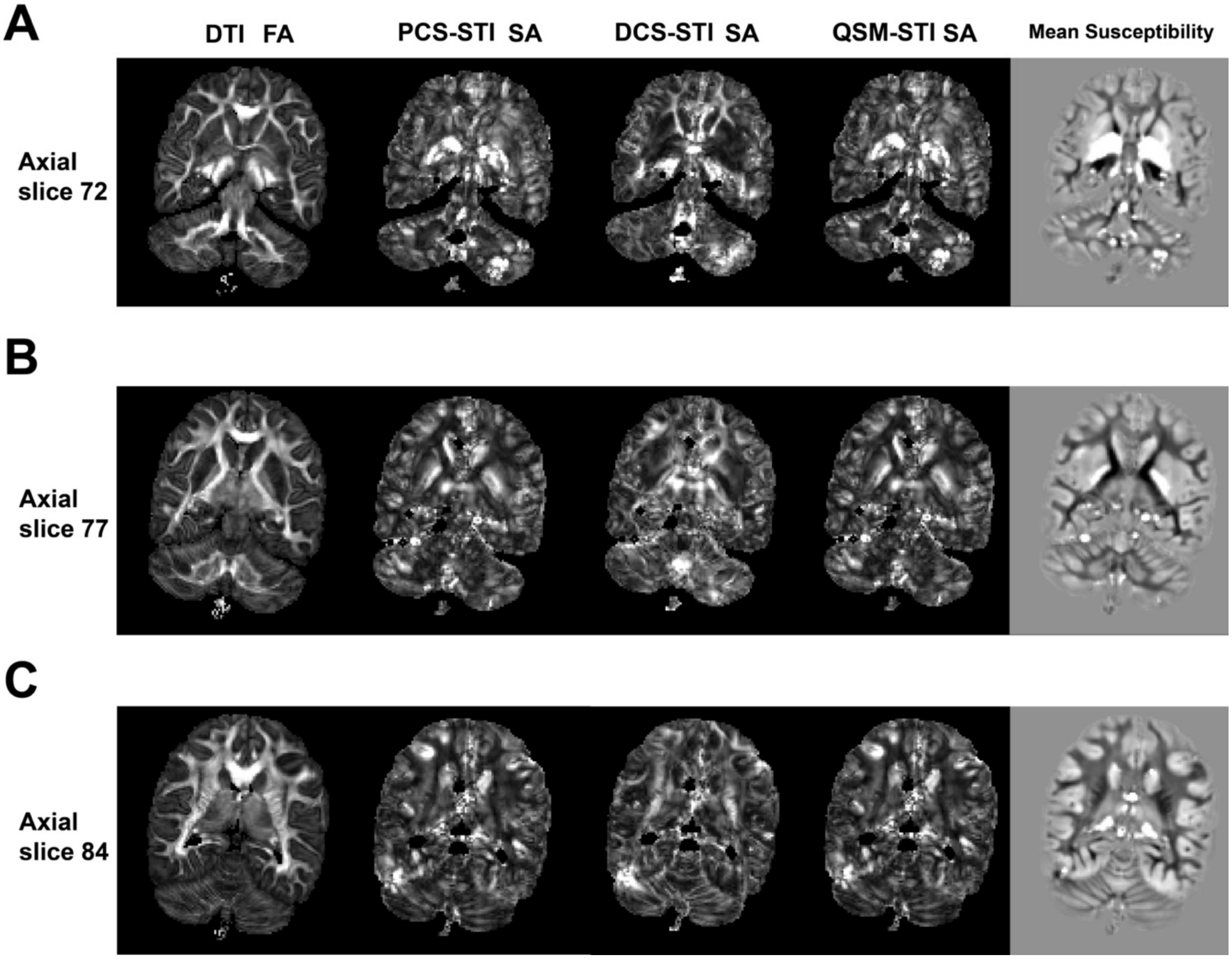

**Supplementary Figure S7**: Simulation scheme for investigating PCS-STI anisotropy. (A) An illustrative fiber covered with iron-rich substances, where Z denotes the fiber's long axis. (B) The cross section of a simulated voxel containing a bundle of fibers covered by iron-rich substances, represented as a 120 x 120 x 200 matrix. The susceptibility for the iron-rich substance is assigned to be 0.3 ppm. (C) The angle between the applied $\mathbf{B}_0$ and the fiber long axis (z) is defined as theta and is sweeping through 0 to 180 degrees. (D) The field map of the middle slice of the simulation when the angle is at 0 degrees (parallel) and 90 degrees (perpendicular). (E) The total field effect is averaged for all the pixels within the simulated voxel. The perturbed field is plotted with respect to the angle between $\mathbf{B}_0$ and the fiber number long axis.

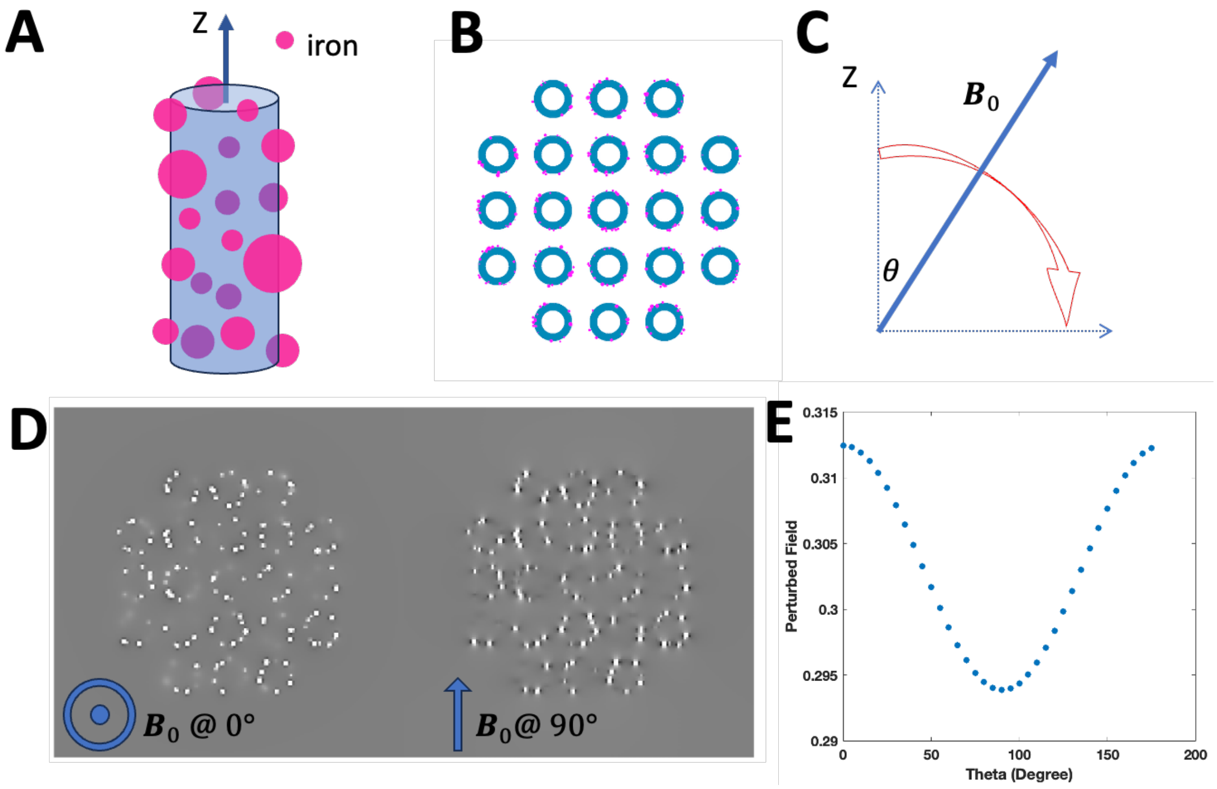
